# More is Different: Constructing the Most Comprehensive Human Protein High-Order Interaction Dataset

**DOI:** 10.1101/2023.11.06.565906

**Authors:** Yuntao Lu, Yongfeng Huang, Tao Li

## Abstract

In biological systems, protein-protein interactions (PPI) weave intricate network patterns that are fundamental to the structural and functional integrity of organisms. While the majority of existing research has been anchored in the study of pairwise PPIs, the realm of high-order interactions remains relatively untapped. This oversight could potentially obscure the deeper intricacies embedded within biological networks. To address this gap, this study formulates a scientific task aimed at predicting high-order protein-protein interactions and introduces a multi-level comprehensive dataset focused on triadic high-order interactions within PPI networks. This dataset incorporates more than 80% of the known human protein interaction relationships and partitions into 60 subsets across a diverse range of functional contexts and confidence. Through meticulous evaluation using cutting-edge high-order network prediction tools and benchmark PPI prediction methodologies, our findings resonate with the principle that “more is different”. Triadic high-order interactions offer a more enriched and detailed informational canvas than their pairwise counterparts, paving the way for a deeper comprehension of the intricate dynamics at play in biological systems. In summary, this research accentuates the critical importance of high-order PPI interactions in biological systems and furnishes invaluable resources for subsequent scholarly investigations. The dataset is poised to catalyze future research endeavors in protein-protein interaction networks, elucidating their pivotal roles in both health and disease states.

## 1. Introduction

Protein-protein interaction (PPI) network models provide a effective approach to investigate the impact of protein interaction patterns on cellular dynamics [1]. By representing protein interaction systems as networks, these models allow for the simulation of the effects of interaction patterns on cellular phenotypes and functions, enabling a deeper understanding of the molecular mechanisms underlying both healthy physiology and diseases [2-4]. While existing PPI research primarily focuses on pairwise relationships between proteins, standard graph models only consider these pairwise interactions, which are represented by the edges connecting two proteins [1,5]. In biology, it is common for multiple proteins to collaborate due to shared functions, thereby forming protein complexes [6-8]. Molecular pathways in cells often involve multi-protein interactions, feedback or feedforward loops and interactions between pathways. Unfortunately, the representation of these pathways of interest is limited by the current reliance on pairwise protein-protein interactions [9-14].

In order to accurately model the intricate relationships in PPI networks, it’s essential to incorporate a higher-order network model. Hypergraphs, which are a generalized representation of networks, offer a more effective way to capture the high-order organization of protein-protein interactions by considering high-order relationships among multiple elements [15-17]. Unlike standard graphs that only account for pairwise connections, hypergraphs extend the model to include relationships with any number of elements. In the context of protein-protein interactions, the “hyperedges” in a hypergraph can represent interactions involving multiple proteins.

However, it is not enough to simply construct high-order networks of protein-protein interactions; it’s equally important to define a clear scientific question that can guide the development of methods to analyze these high-order network data and gain biological insights into cellular functions and phenotypes. Triadic protein-protein interactions, which involve three proteins, are particularly significant as they serve as a gateway to studying higher-order protein relationships. They are not merely an extension of pairwise; they represent a crucial transition from low-order to high-order interaction studies. By thoroughly investigating triadic protein-protein interaction, we can not only uncover previously unknown information in the protein network, but also pave the way for future research on more complex multi-body protein responses.

The primary objective of this study is to formulate the scientific task of predicting high-order protein-protein interactions, with a specific focus on triadic interactions among proteins. To accomplish this, this study has assembled a dataset of triadic protein-protein interactions and partitioned into 60 subsets focusing primarily on the confidence associated with triadic relationships and the specific functions of the proteins involved. Our study undertakes a thorough evaluation of various hyperedge prediction methods, shedding light on the profound implications of high-order relationships in protein-protein interaction prediction. We underscore the “more is different,” highlighting the intricate and multifaceted nature of protein-protein interactions.

## 2. Task Formulation

### 2.1 Hypergraph Fundamentals

Hypergraphs serve as an extension to conventional graph models by accommodating relationships involving an arbitrary number of elements, thereby generalizing the concept of strictly pairwise edges found in traditional graphs. To elucidate the distinctions between low-order and high-order network representations, consider an illustrative example. Assume there are seven sets of protein-protein interactions with the following protein compositions: [A, B, C, D], [E, F, G], [A, B], [A, H], [B, H], [D, E], [G, H]. In a conventional network framework, the triad of relationships [A, B], [B, H], [A, H] would be represented as a triangle. However, the triad [E, F, G] would also be depicted as a triangle, leading to ambiguity. This inherent limitation of low-order networks is effectively addressed in high-order networks, where the two scenarios can be distinctly represented, as illustrated in **Figure 1a**.

**Figure 1.**
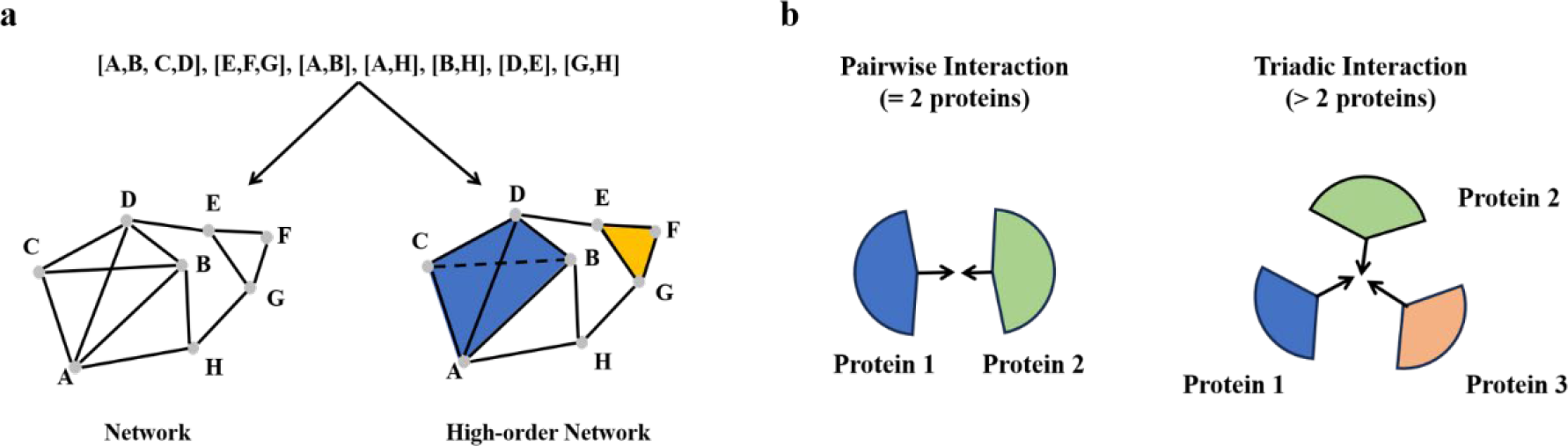
Visualization of Triadic Relationships in Protein-Protein Interactions. **(a) Hypergraphs Explained:** A conceptual illustration showcasing the structure of hypergraphs. High-order networks adeptly overcome the constraints observed in traditional low-order networks. **(b) From Pairs to Triads:** A schematic contrasting the simple pairwise interactions with the more complex triadic interactions in protein networks.

### 2.2 Hypergraph Representation in Triadic Protein-Protein Interactions

In the realm of triadic protein-protein interactions, a “hyperedge” within the hypergraph framework signifies high-order interactions involving three proteins. Specifically, a protein-protein interaction network is denoted as *H* = {*V, E_h_*}, where *V* = {*v*_1_, *v*_2_, … *v*_*n*_} represents proteins and *E*_*h*_ = {*e*_1_, *e*_2_ …. *e*_*m*_}, with *e*_*p*_ ∈ *V* (*p* = 1,2 … *m*) *and* |*e*_*p*_| = 3 signifies triadic protein-protein interactions (**Figure 1b**). If two nodes share the same hyperedge, they are termed as adjacent nodes. The association matrix of the hypergraph is denoted by *H* ∈ *R*^*n*×*m*^, which consists of logical values indicating the relationship between a given node and the hyperedge. Specifically, If node v_i_is part of e_p_, then *H*(*i,p*) = 1, otherwise, it is 0.

### 2.3 Prediction Objectives and Functions

The central aim of triadic protein-protein interaction prediction is to identify the most probable hyperedges within the observed hyperedge set *ε*. To realize this, the majority of hyperedge prediction methodologies strive to learn a function Ψ, which assigns a score to a given potential hyperedge *e*, thereby aiding in its validation. The function Ψ(e) is defined as follows:

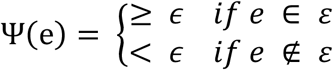

where *ϵ* serves as a threshold to convert the continuous value into a discrete label.

## 3. Dataset Construction

The dataset curated in this study is a comprehensive compilation of human triadic protein-protein interactions. It has been meticulously structured to provide a multi-dimensional perspective, focusing primarily on the confidence associated with tririadic relationships and the specific functions of the proteins involved (**Figure 2**).

**Figure 2.**
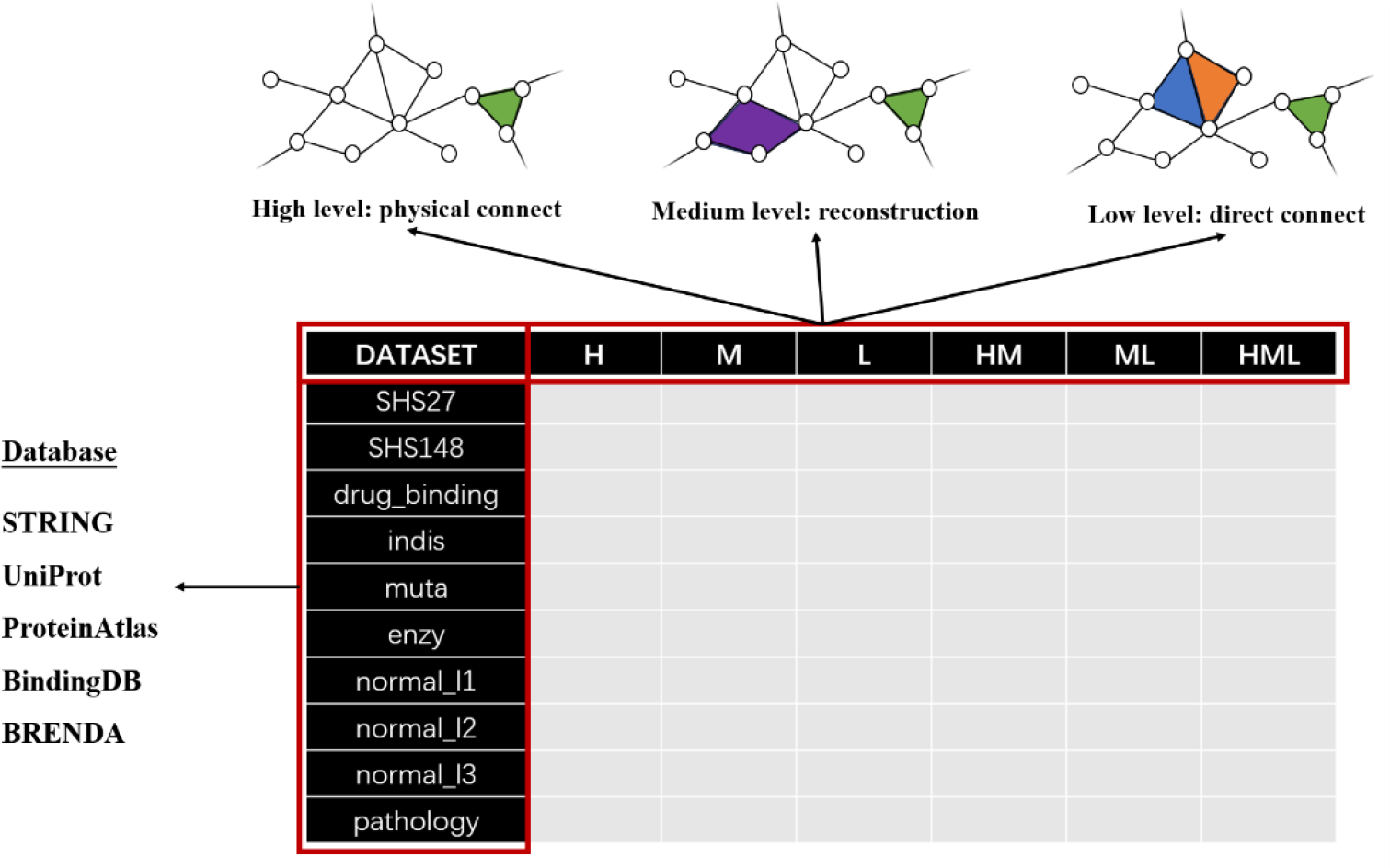
Overview of the Dataset’s Structure. This figure delineates the bifurcated structure of the dataset. Vertically, subsets are categorized based on the confidence levels associated with the triadic relationships. Horizontally, they are segmented according to the specific functional roles of the encompassed proteins. Together, this multi-dimensional dataset incorporates 60 subsets and offers a comprehensive and nuanced view of PPI, serving as a treasure trove for researchers delving into human protein-protein interaction intricacies.

From a confidence standpoint, the triadic relationships is categorized into three distinct levels: high(H), medium(M) and low(L). Based on these confidence levels, six combination subsets are derived, namely H, M, L, HM, ML and HML. This stratification ensures that researchers can select subsets based on the reliability of the interactions, allowing for more targeted and accurate analyses.

From a functional standpoint, the dataset delves deeper by constructing ten functional subsets. These subsets are derived by integrating information from two public datasets, SHS27 and SHS148, and by considering various protein functional preferences. Such a functional breakdown ensures that the dataset not only captures the interactions but also provides insights into the specific roles and implications of these interactions in biological processes.

Consequently, the dataset is partitioned into a total of 60 subsets, each offering a unique perspective on triadic protein-protein interactions. This vast and diverse dataset encapsulates a wealth of information, making it an invaluable resource for researchers aiming to unravel the complexities of protein-protein interactions in human biology.

### 3.1 Database Source

The dataset curated in this study is primarily composed of human protein pairwise interaction data, a plethora of protein functional information and two publicly available datasets.

#### Pairwise Interaction Data Collection

The foundational data for this dataset, which pertains to human protein-protein interactions, was meticulously extracted from the STRING database (https://string-db.org/).To ensure the reliability and accuracy of the interactions, only those with a score surpassing 700 were incorporated into the dataset.

#### Acquisition of Protein Functional Information

A comprehensive understanding of protein functions is pivotal for any protein-protein interaction dataset. To this end, the study sourced protein functional data from the UniProt database (https://www.uniprot.org/). Additionally, data pertaining to the functional roles of proteins in human tissues was derived from the ProteinAtlas database (https://www.proteinatlas.org/). To further enhance the dataset’s depth, information regarding protein-drug interactions was incorporated from the BindingDB database (https://www.bindingdb.org/). Lastly, to provide insights into enzymatic functions, enzyme-related data was retrieved from the BRENDA database (https://www.brenda-enzymes.org/).

#### Public Dataset Integration

To ensure a holistic and comprehensive dataset, external datasets were also integrated. Specifically, the SHS27 and SHS148 datasets, which are pivotal in the realm of protein-protein interactions, were sourced from an open-access repository (https://github.com/muhaochen/seq_ppi) [18].

### 3.2 Categorization Based on Confidence Levels

To uphold the highest standards of rigor and accuracy, three distinct confidence levels for triadic protein-protein interactions were established. The low confidence (L) is primarily predicated on pairwise high-confidence interactions among the three proteins, resulting in open triangular structures within the network. The medium confidence (M) leverages a validated Bayesian approach to pinpoint network segments that can be attributed to potential high-order interactions. Given that this method may identify high-order relationships involving more than three proteins, this study made a clear distinction between triadic and multi-body relationships within the dataset. The high confidence (H) is solely reliant on protein spatial configurations. When three proteins form a physical complex, it is deemed a highly reliable triadic relationship. Based on these confidence tiers, the dataset was further segmented into six subsets (H, M, L, HM, ML, HML), thereby providing a robust foundation for future subsequent multigranularity studies.

A graph *G* = {*V, E*} is defined, where *V* symbolizes proteins and *E* denotes pairwise interactions. Within this framework, triadic protein represents high-order relationships involving the collective interaction of three proteins.

#### Low confidence

A set *V*_h_ = (*v*_*a*_, *v*_*b*_, *v*_*c*_) is classified as a triadic protein-protein interaction if (*v*_*a*_, *v*_*b*_), (*v*_*a*_, *v*_*c*_), and (*v*_*b*_, *v*_*c*_) are all elements of *E*.

#### Medium confidence

High-order information are inferred from low-order networks using a methodology proposed by Young et al. [19]. This involves the construction of a Bayesian generative model that explicates how the network data *G* is generated in the presence of potential high-order interactions *H*. This model is designed to estimates the posterior probability *P*(*H*|*G*) based on the observed network *G*.

#### High confidence

For a subgraph *G*_*ph*_ = (*V*_*ph*_, *E*_*ph*_) with *G*_*ph*_ ∈ *G*, a set 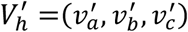 is deemed a triadic protein-protein interaction if 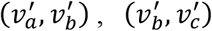 and 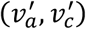 all belong to *E*_*ph*_.

### 3.3 Categorization Based on Protein Functionality

To ensure a comprehensive understanding of protein functions and their implications in various biological contexts, this study meticulously categorized proteins based on distinct functional perspectives. The objective was to ensure diversity across functional datasets while maintaining a consistent volume of data within each category.

#### Drug binding Proteins Set

This category focuses on proteins that play pivotal roles in drug interactions. Candidate proteins that function as potential drug targets and exhibit interactions with small-molecule ligands are identified through the BindingDB database. These proteins are subsequently labeled as ‘drug_binding proteins’.

#### Disease-Associated Proteins Set

Understanding disease-associated proteins is crucial for therapeutic and diagnostic advancements. Proteins that have associations with diseases due to genetic variations are sourced from the UniProt database. These are labeled as ‘indis proteins’.

#### Enzymatic Proteins Set

Enzymes play a central role in various biochemical reactions. Proteins that are classified as functional enzymes are extracted from the BRENDA database and are labeled as ‘enzy proteins’.

#### Mutated Proteins Set

Mutations can drastically alter protein functions, often leading to diseases. Proteins that experience changes in their biological properties due to experimental mutations in one or more amino acids are identified using the UniProt database. These are labeled as ‘muta proteins’.

#### Normal Tissues Proteins Set (Levels1-3)

The expression of proteins in normal tissues provides insights into their physiological roles. Based on the ProteinAtlas database, proteins are categorized into three levels based on the reliability of their expression profiles in human tissues, as determined through immunohistochemistry. These proteins are labeled as ‘normal_(l1-l3) proteins’.

#### Pathological Proteins Set

Pathological proteins often have implications in disease progression and patient outcomes. Using the ProteinAtlas database, proteins are identified based on their staining patterns in human tumor tissues through immunohistochemistry. Proteins that correlate strongly with patient survival rates and exhibit high staining levels in a majority of patient samples are labeled as ‘pathology proteins’.

## 4. Analyzing the Dataset

### 4.1 Crafting a Comprehensive Multi-Level Dataset for Human Triadic Protein-Protein Interactions

In an effort to create the most comprehensive multi-level human triadic protein-protein interaction dataset, this study amalgamated two public datasets (SHS27, SHS148), eight functional protein datasets and an exhaustive human protein interaction dataset. This integration resulted in the successful construction of ten triadic protein-protein interaction datasets of varying scales (**Table 1**). Notably, the eight functional protein datasets exhibited a substantial increase in both the number of proteins and PPIs when compared to SHS27(**Figure 3a**). Collectively, these functional protein datasets encompass 79.5% of proteins and 84% of PPIs in human protein-protein interactions, underscoring the dataset’s comprehensiveness and broad applicability.

**Table 1:** Dataset statistics, respectively describing the number of proteins in the dataset, the number and density of PPIs.

| Dataset | #Proteins | #PPIs | #Edge density |
| --- | --- | --- | --- |
| SHS27 | 1553 | 6660 | 0.0055 |
| SHS148 | 5189 | 44488 | 0.0033 |
| drug_binding | 3129 | 71574 | 0.0146 |
| indis | 4817 | 99153 | 0.00855 |
| muta | 4885 | 104590 | 0.00877 |
| enzy | 2538 | 38939 | 0.0124 |
| normal_l1 | 4051 | 38013 | 0.00463 |
| normal_l2 | 1285 | 18046 | 0.0219 |
| normal_l3 | 3347 | 34514 | 0.00656 |
| pathology | 4779 | 79527 | 0.00697 |
| total | 16814 | 252984 | 0.00179 |

**Figure 3.**
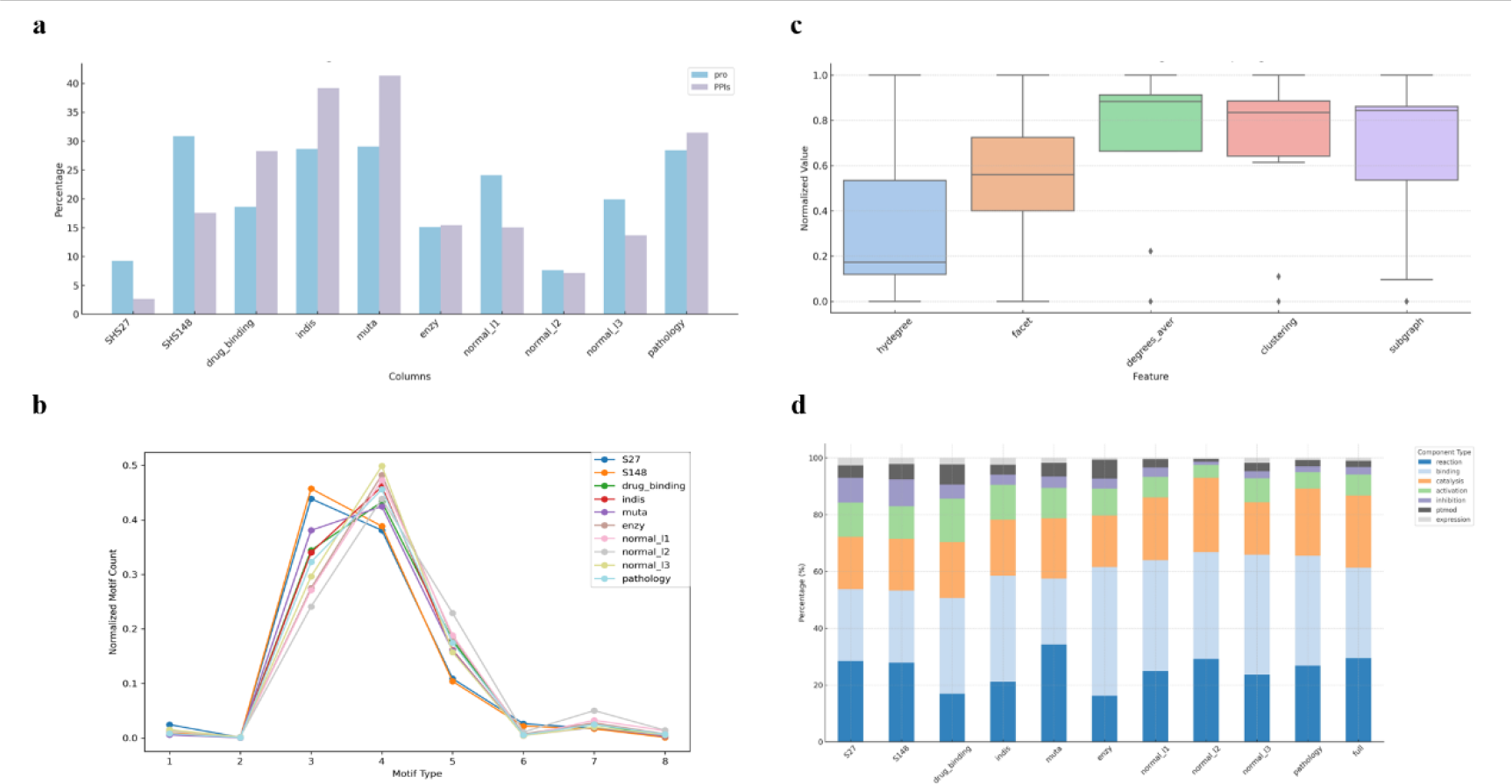
Insights from the Multi-level Human Triadic Protein-Protein Interaction Dataset. **(a) Coverage Diversity:** A depiction of protein and PPI coverage across the diverse datasets. The variations in protein and pairwise interaction coverage across datasets provide a lens into the unique information content inherent to each. **(b) Consistent Motif Patterns:** Motif distribution curves for each dataset, emphasizing the striking consistency across various functional subnetworks of human protein-protein interactions from the low-order perspective. **(c) High-order Topological Differences:** A comparative analysis showcasing the differences between low-order and high-order topological features derived from each dataset. This contrast underscores ‘more is different’, the enhanced depth and richness of information offered by high-order networks. **(d) Interaction Type Spectrum:** A breakdown of the types of PPI interactions present within each dataset. While each dataset consistently showcases certain biological features, clear functional distinctions are also evident, highlighting the dataset’s diversity.

The low-confidence subset encapsulates all conceivable triadic protein-protein interactions derived from established high-confidence protein-protein interactions. The high-confidence subset embodies the most reliable segment of potential triadic relationships. Medium confidence serves as a transitional phase, bridging latent triadic interactions to dependable higher-order relationships through advanced network theory. The distribution of data volumes across six subsets within the ten datasets is delineated in **Table 2**, affording researchers a versatile array of options to cater to diverse research requirements.

**Table 2:**
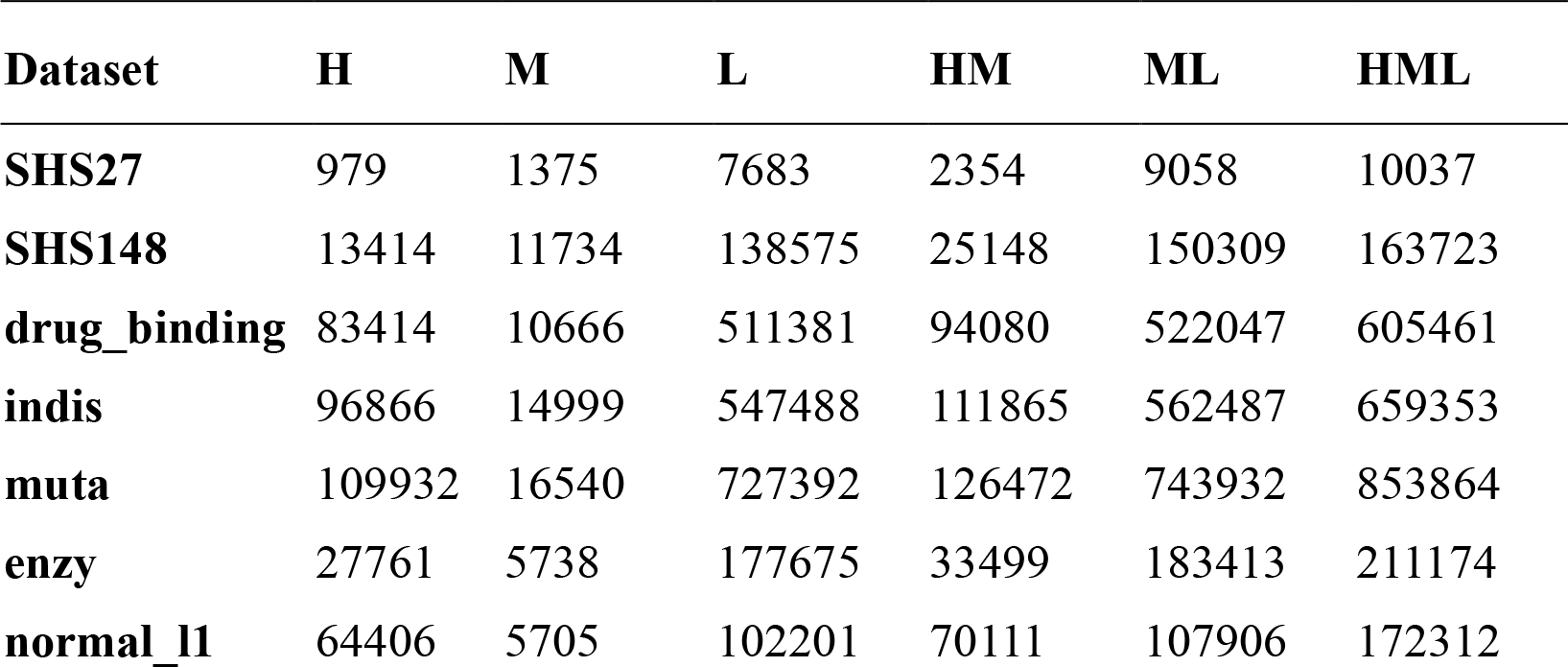
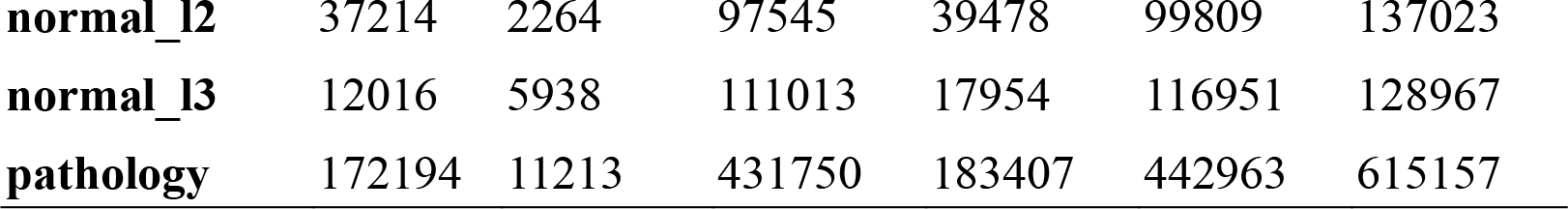
Statistics of the triadic protein-protein interactions at 60 subsets.

| Dataset | H | M | L | HM | ML | HML |
| --- | --- | --- | --- | --- | --- | --- |
| SHS27 | 979 | 1375 | 7683 | 2354 | 9058 | 10037 |
| SHS148 | 13414 | 11734 | 138575 | 25148 | 150309 | 163723 |
| drug_binding | 83414 | 10666 | 511381 | 94080 | 522047 | 605461 |
| indis | 96866 | 14999 | 547488 | 111865 | 562487 | 659353 |
| muta | 109932 | 16540 | 727392 | 126472 | 743932 | 853864 |
| enzy | 27761 | 5738 | 177675 | 33499 | 183413 | 211174 |
| normal_l1 | 64406 | 5705 | 102201 | 70111 | 107906 | 172312 |
| **normal_l2** | 37214 | 2264 | 97545 | 39478 | 99809 | 137023 |
| **normal_l3** | 12016 | 5938 | 111013 | 17954 | 116951 | 128967 |
| **pathology** | 172194 | 11213 | 431750 | 183407 | 442963 | 615157 |

### 4.2 Gleaning Topological Insights from High-Order Interactions

To gain more nuanced insights into protein-protein interaction (PPI) networks, this study conducted an evaluation of the topological features inherent to each PPI network. These features predominantly capture the connectivity structures within the network, which are intrinsically linked to the interaction patterns of proteins at the cellular level. Analysis of the eight functional datasets disclosed consistent topological characteristics in low-order network features, particularly in the distribution curves of network motifs. Various functional subnetworks within human protein interactions exhibited remarkable uniformity (**Figure 3b**). However, a notable divergence was observed in high-order features across these datasets (**Figure 3c**), accentuating that ‘more is different’, the depth advantage conferred by high-order networks in information extraction.

### 4.3 Assessing Coverage and Diversity in Triadic Protein-Protein Interactions

Triadic protein-protein interactions account for approximately 50% of the total human protein interactions, albeit with variations across different datasets. This observation underscores that high-order protein-protein interactions are both ubiquitous and varied. High coverage ensures a wealth of information within the dataset, thereby facilitating small-sample learning tasks and offering a comprehensive view of human protein interaction patterns. The diversity in interactions among functionally distinct proteins is pivotal for probing into the intricacies of protein functions and biological processes.

The datasets assembled in this study serve as a testament to the diversity and complexity inherent in triadic protein-protein interactions. Each dataset consistently manifests biological features, notably significant proportions of binding and reaction events (**Figure 3d**). However, discernible functional disparities are also evident among the datasets, potentially attributed to variations in data sources, collection methodologies or biological backgrounds. These datasets not only furnish researchers with a profound understanding of the types of protein-protein interactions but also encapsulate the differences and consistencies in these interactions across diverse biological conditions.

## 5. Experiments

### 5.1 Evaluating Contemporary Hypergraph Prediction Methods for the Triadic Protein-Protein Interaction Task

#### 5.1.1 An Overview of Selected Prediction Methods

Various computational approaches have been devised to predict high-order relationships within networks. These approaches can be broadly classified into four distinct categories: similarity-based, probability-based, matrix optimization-based and deep learning-based [20]. To rigorously assess their efficacy on the triadic protein-protein interaction dataset, we handpicked representative algorithms from each category for an exhaustive study.

**HPRA** [21]: The similarity-based prediction method leverages shared neighbors and the Jaccard coefficient as metrics to evaluate node similarity and predict potential hyperedges.

**HPLSF** [22]: This probability-based method employs the adjacency and degree matrices of the hypergraph to gauge node similarity, also incorporating shared neighbors and the Jaccard coefficient.

**C3MM** [23]: This matrix optimization-based method predicts hyperedges by capturing node similarity through the adjacency and degree matrices of the hypergraph.

**HyperSGCN** [24] & **NHP** [25]: These deep learning-based method utilize graph/hypergraph-centric neural network architectures to capture high-order topological features, thereby substantially boosting the performance of hyperedge prediction.

#### 5.1.2 Metrics and Outcomes of the Evaluation

In the evaluation process, Area Under the Precision-Recall Curve (AUPR) and F1 Score were employed as the key performance indicators. After scrutinizing the high-quality triadic interaction subset (H set) across all functional datasets (**Figure 4, Supplementary Table 1**), it was discerned that the deep learning-based HyperSGCN algorithm outperformed its counterparts in overall performance. It’s noteworthy that the predictive outcomes varied across different functional datasets for all methods. This variation underscores the distinct high-order informational content inherent in each functional protein dataset, highlighting the intrinsic heterogeneity among subsets.

**Figure 4.**
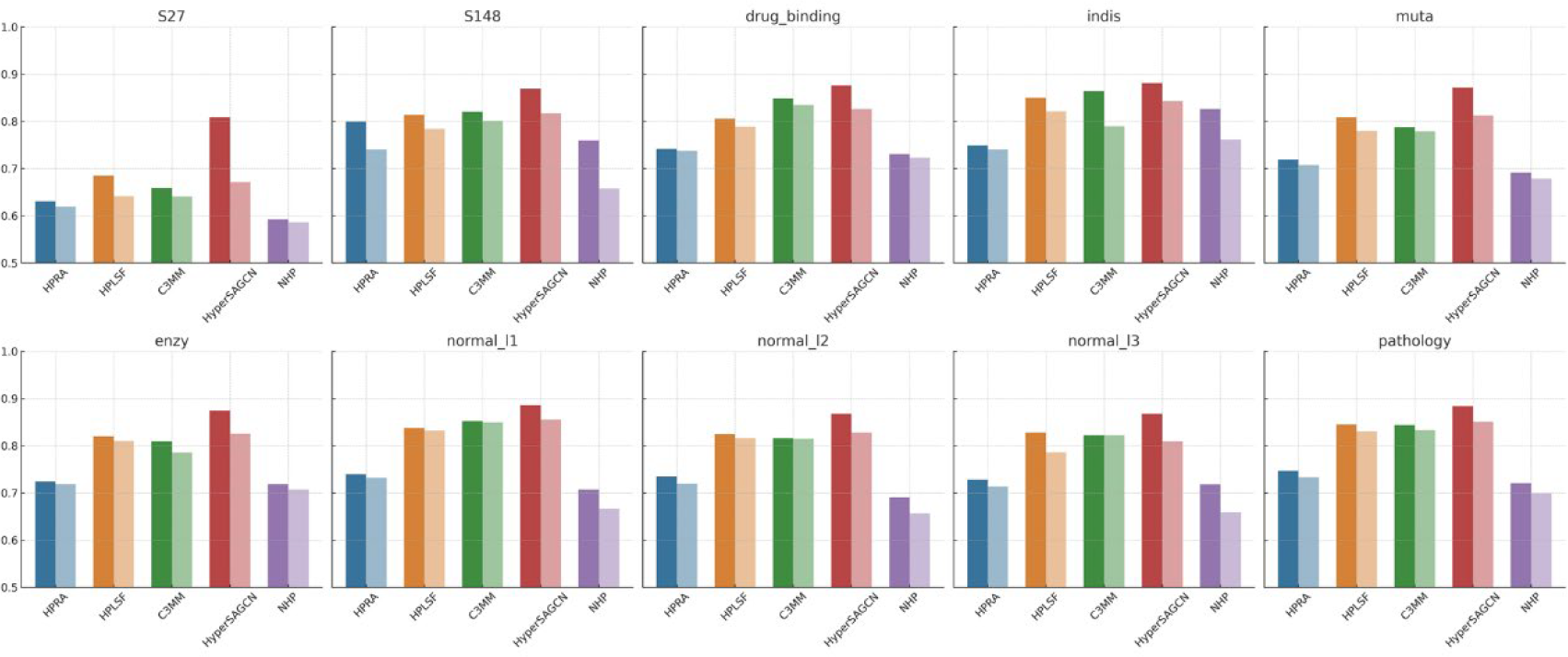
Evaluating Hypergraph Prediction Methods. A performance comparison of various hypergraph prediction methods, with the left side representing F1 score and the right side showcasing AUPR across different datasets. It’s evident that the efficacy of each algorithm varies across functional subsets. Among the contenders, HyperSAGCN distinguishes itself by consistently delivering superior results across all datasets, reinforcing its prowess in accurately predicting hypergraph interactions.

### 5.2 In-depth Analysis of Dataset’s Informative Capacity

In an effort to further understand the depth and breadth of our dataset’s informativeness, this study embarked on an extended evaluation. Utilizing the HyperSGCN algorithm, an exhaustive analysis was conducted across all datasets, spanning multiple levels of granularity.

#### 5.2.1 Consistency in Performance Across Datasets

An in-depth evaluation of HyperSGCN across various datasets was conducted. Results presented in **Supplementary Table 2** and **Figure 5a** indicate that the model maintains consistent performance across all functional protein datasets, especially in the high-confidence (H) subset. Interestingly, an increase in the volume of subset data did not correspond to an improvement in prediction accuracy. This observation intimates a significant level of information heterogeneity across datasets with varying confidence levels, thereby unveiling distinct layers of protein-protein interactions and providing valuable insights into their intricate relationships.

**Figure 5.**
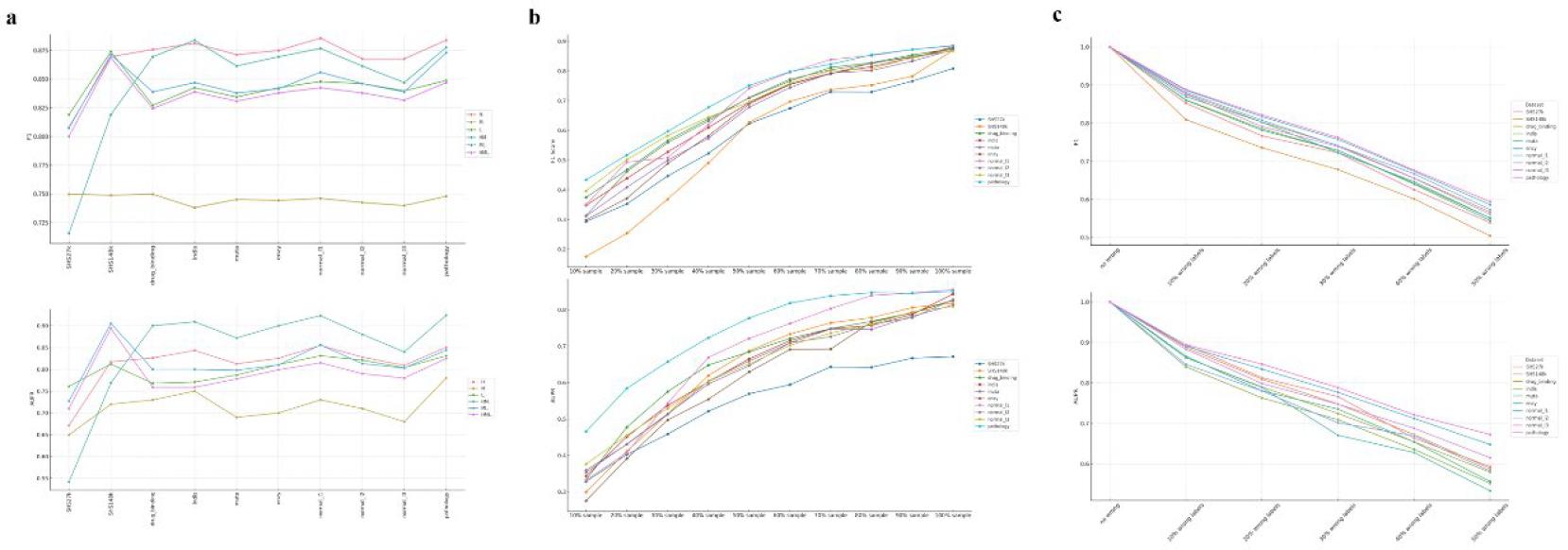
In-depth Analysis of Dataset Informativeness with HyperSAGCN. **(a) Consistency Across Datasets:** A depiction of performance consistency across all subsets. The uniform performance, especially within the high-confidence (H) subset, suggests a notable level of information heterogeneity across datasets with different confidence levels. **(b) Saturation Test:** Performance evaluations based on retaining 10%-100% of training samples in the H subset of each function dataset, spread across ten gradients. The outcomes highlight the dataset’s minimal redundancy, ensuring its richness and diversity. **(c) Robustness Test:** Performance evaluations after introducing 10%-50% erroneous samples into the training set, spanning five gradients. The findings suggest that while the dataset is comprehensive, current state-of-the-art methods may not exhibit optimal resilience when applied to it. For all panels, the top graph delineates the F1 score, while the bottom graph presents the AUPR score.

#### 5.2.2 Saturation Analysis of Information

To probe the dataset’s information redundancy, an information saturation test was executed on the high-confidence (H) subset across all function datasets (**Figure 5b, Supplementary Table 3**). In this saturation test, the predictive performance was assessed by retaining 10%-100% of the training samples in the H subset of each dataset, evaluated across ten gradients. Interestingly, as the data volume escalates, the performance curve exhibits a gradual decay in its curvature, indicating that the dataset possesses a low degree of redundancy.

**Table 3:**
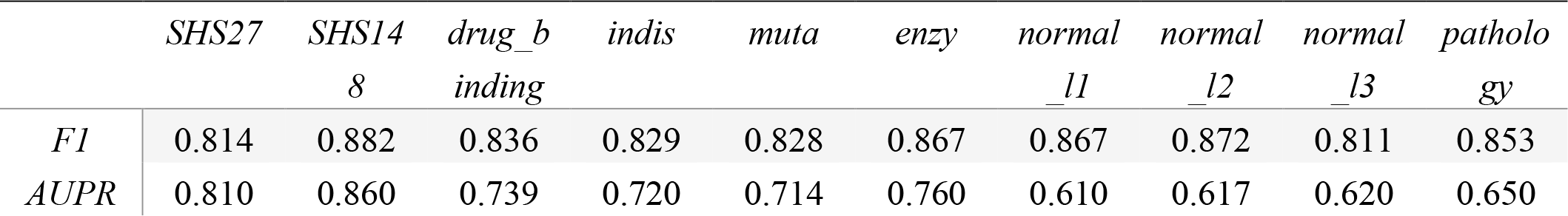
Performance of HIGHPPI in predicting protein-protein interactions across various function datasets.

#### 5.2.3 Testing Dataset Robustness

To evaluate the dataset’s resilience to perturbations, a robustness test was initiated on the high-confidence (H) subset across all function datasets (**Figure 5c, Supplementary Table 4**). In this test, the predictive performance was evaluated by introducing 10%- 50% erroneous samples into the training set, again evaluated across five gradients. Notably, as the proportion of erroneous information escalated, the prediction performance exhibited a gradual decline. This observation implies that the current state-of-the-art methods do not demonstrate optimal robustness when applied to this dataset. Such a trend emphasizes the need for developing more resilient prediction methods on the triadic interaction task, ensuring consistent performance even in the presence of perturbations.

### 5.3 More is different: Exploring the Impact of High-Order Relationships on PPI Prediction

#### 5.3.1 Benchmark Analysis Using HIGHPPI

HIGHPPI [26] is a cutting-edge algorithm that amalgamates both sequence and structural information of proteins, utilizing hierarchical graphs [27] for the prediction of PPI relationships. Among the plethora of existing PPI prediction algorithms, HIGHPPI distinguishes itself in terms of performance. Therefore, this study elected HIGHPPI as a benchmark to explore the potential benefits of incorporating high-order features to augment PPI prediction accuracy.

Initial evaluations were conducted across diverse datasets using HIGHPPI to establish a foundational baseline for PPI prediction. As delineated in **Table 3**, performance of HIGHPPI fluctuates across datasets, likely attributable to the unique characteristics and inherent complexities of each dataset.

#### 5.3.2 The Significance of High-Order Features in Predictions

To probe further into the impact of high-order features, the DUNE [28] algorithm was employed to extract hypergraph representations from the hypergraph network, subsequently integrating them into the feature matrix. Encouragingly, as depicted in **Figure 6 and Supplementary Table 5**, the incorporation of high-order information consistently elevated prediction accuracy across the majority of datasets. This implies that high-order features endow the model with a rich contextual framework, thereby facilitating more precise predictions of PPIs. This finding corroborates our preliminary hypothesis that protein-protein interactions are not merely binary but often encompass intricate multi-entity interactions.

**Figure 6.**
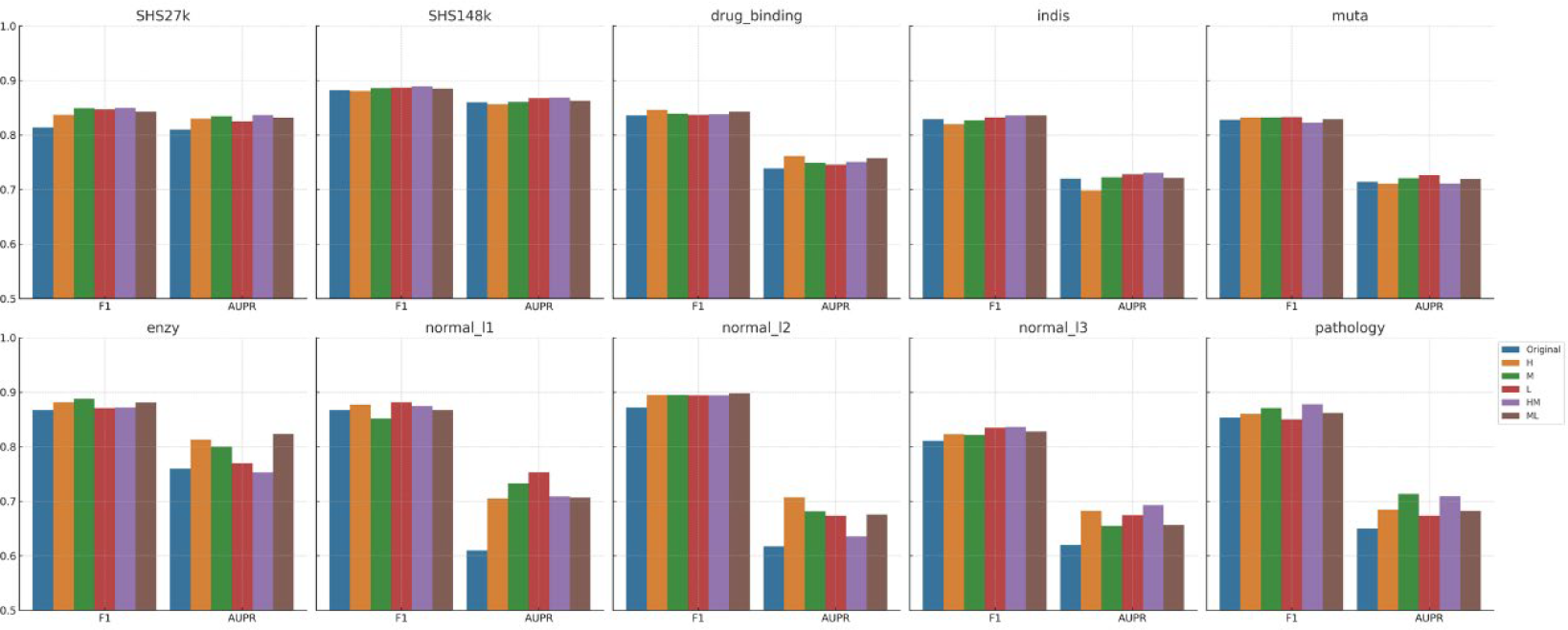
Leveraging High-Order Features for Enhanced PPI Predictions. Performance improvements in PPI predictions across various datasets are showcased when high-order features from triadic protein-protein interactions are integrated using HIGHPPI. The consistent enhancement in prediction accuracy across most datasets underscores the value of high-order information. This suggests that these features provide the model with a rich contextual backdrop, enabling more accurate and nuanced predictions of protein-protein interactions.

Interestingly, the degree of prediction enhancement differed across function datasets when various subsets were employed. Overall, the HM subset manifested the most significant uplift in prediction outcomes. This diverges from the performance trends observed in triadic relationship predictions and is likely because the HM subset harbors a greater wealth of high-order information, which proves instrumental in PPI prediction. This further underscores the pivotal role of high-order information in PPI prediction. Such insights offer a renewed perspective on the intricate network architecture of proteins, thereby serving as a valuable compass for future research endeavors.

## 6. Discussion

This study successfully formulated a scientific task aimed at predicting high-order protein-protein interactions and engineered a comprehensive, multi-level triadic protein-protein interaction dataset and rigorously evaluated a variety of hyperedge prediction methodologies. While the study represents a meaningful advancement in the field, it also acknowledges certain limitations in this study, which in turn delineate avenues for future research endeavors.

### 6.1 The Unexplored Dimension: Temporal Dynamics

One significant limitation in the current study is the absence of temporal dynamics. Existing static network models are inadequate for capturing dynamic transitions, such as those between open and closed triangles or from the L subset to the H subset. Given the protein-protein interactions are inherently dynamic entities, evolving in response to time and environmental changes, future research should aim to integrate temporal dimensions into the modeling framework. This might necessitate the acquisition of time-series data or the development of novel models tailored for dynamic high-order networks.

### 6.2 Venturing Beyond Mere Triadic Interactions

Future prediction tasks should extend beyond the realm of mere triadic protein-protein interactions. In biological systems, complex interactions frequently involve multiple proteins, potentially including quartets, quintets or even larger assemblies. An emergent research direction should thus focus on the creation of new models and algorithms capable of addressing these intricate multi-entity interactions. This will likely require the gathering of additional high-order protein-protein interaction data or the invention of innovative data processing techniques.

### 6.3 Broadening the Dataset’s Horizon

There is an aspiration to broaden the dataset to include a wider array of biological species, thereby enhancing its diversity and representational scope. Such an expansion would enable more comprehensive downstream tasks and deepen our understanding of the variances and commonalities across different species.

### 6.4 Challenges in Algorithmic Efficiency and Complexity

Current high-order network algorithms are not without their drawbacks. Particularly when grappling with large-scale networks, many extant open-source algorithms demonstrate suboptimal performance, often coupled with elevated time complexities. This highlights an imperative for future research to concentrate on algorithmic optimization, especially in the context of large-scale and complex networks, to improve both efficiency and accuracy.

## 7. Conclusion

This study represents a seminal contribution to the field of Protein-Protein Interaction (PPI) research by introducing a comprehensive, multi-level dataset focused on triadic high-order interactions. The dataset was rigorously evaluated using state-of-the-art high-order network prediction algorithms and PPI forecasting methods. The findings unequivocally demonstrate that ‘more is different’, high-order interactions, particularly triadic interactions, offer a richer informational context than traditional pairwise interactions. This enables a more nuanced understanding of the complex mechanisms governing biological systems.

## Data Availability

The dataset supporting the conclusions of this article is available for academic and research purposes. It encompasses a comprehensive collection of human triadic protein-protein interactions, amalgamated from multiple sources, including public datasets and functional protein datasets. The dataset provides a detailed view of high-order interactions within protein-protein interaction networks, making it an invaluable resource for researchers aiming to delve deeper into the realm of protein-protein interaction prediction. Detailed statistics, topological insights and other relevant visualizations are also included to aid in the understanding and utilization of the dataset. For access, researchers can refer to the repository link (https://github.com/laxlyt/triadic-protein-interactions/) provided in the article or contact the corresponding author.

## Fund

This research was supported by the National Natural Science Foundation of China (Grant No. 92251304), and the National Key Research and Development Program of China (Grant No. 2020YFA0907402 and No. 2018YFA0903100).

